# Knockout studies using CD34+ hematopoietic cells suggest that CD44 is a physiological human neutrophil E-selectin ligand

**DOI:** 10.1101/2023.08.18.553923

**Authors:** Yuqi Zhu, Sriram Neelamegham

## Abstract

The recruitment of peripheral blood neutrophils at sites of inflammation involves a multistep cascade, starting with E- and P-selectin expressed on the inflamed vascular endothelium binding sialofucosylated glycans on leukocytes. As the glycoconjugate biosynthesis pathways in different cells are distinct, the precise carbohydrate ligands of selectins varies both across species, and between different immune cell populations in a given species. To study this aspect in human neutrophils, we developed a protocol to perform CRISPR/Cas9 gene-editing on CD34+ hHSCs (human hematopoietic stem/progenitor cells) as they are differentiated towards neutrophil lineage. This protocol initially uses a cocktail of SCF (stem-cell factor), IL-3 (interleukin-3) and FLT-3L (FMS-like tyrosine kinase 3 ligand) to expand the stem/progenitor cells followed by directed differentiation to neutrophils using G-CSF (granulocyte colony-stimulating factor). Microfluidics based assays were performed on a confocal microscope platform to characterize the rolling phenotype of each edited cell type in mixed populations. These studies demonstrated that CD44, but not CD43, is a major E-selectin ligand on human neutrophils. The loss of function results were validated by developing sialofucosylated recombinant CD44. This glycosylated protein supported both robust E-selectin binding in a cell-free assay, and it competitively blocked neutrophil adhesion to E-selectin on inflamed endothelial cells. Together, the study establishes important methods to study human neutrophil biology and determines that sialoflucosylated-CD44 is a physiological human E-selectin ligand.

**KEY POINTS:**

- A CRISPR-Cas9 based protocol was optimized to knockout genes in primary human neutrophils derived from CD34+ hHSCs
- Under physiological fluid shear conditions, a sialofucosylated form of CD44 expressed on human neutrophils is a major E-selectin ligand

**VISUAL ABSTRACT:** 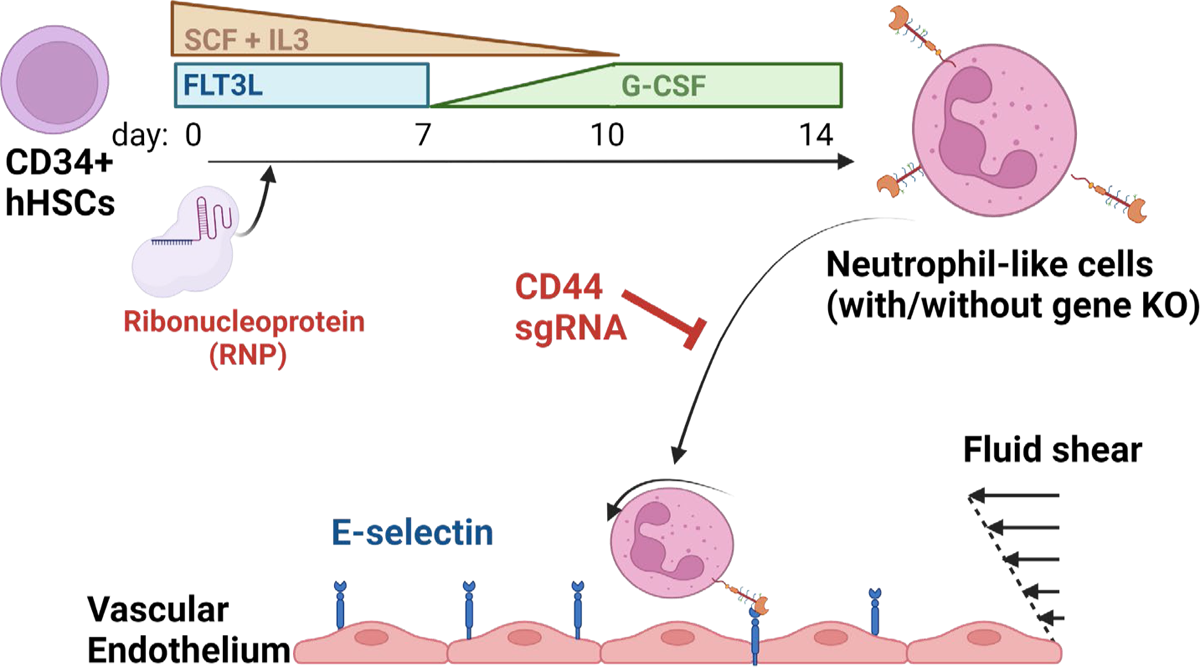

## INTRODUCTION

Neutrophils are short-lived blood cells with lifetimes of 19±6 hours in circulation^1^. In the healthy human population, they constitute 50-70% of circulating leukocytes and act as the first line of defense to combat infection. Besides their well-known role in acute inflammatory response, recent findings suggest that neutrophils also contribute to chronic inflammatory conditions like atherosclerosis and autoimmune diseases^2^. This expanded role of neutrophils is in part due to the realization that neutrophils can engage lymphocyte and antigen-presenting cells, both at sites of inflammation and in the draining lymph nodes, and by their presentation of a variety of soluble and insoluble factors^3^. To add to this complexity, data continue to accumulate suggesting that neutrophils may not be a homogenous population of short-lived cells^4,5^. Additionally, it is now recognized that human Neutrophil to Lymphocyte Ratio (NLR) increases during cancer^6^, and tumor-associated neutrophils may promote cancerous growth and metastasis^7–10^. These observations suggest the need to develop new tools to better understand neutrophil biology.

Except for small vessels of the liver^5^ and lungs^11^, the accumulation of neutrophils in tissue is typically initiated by protein-carbohydrate interactions involving the selectin family of adhesion molecules. Three members of the selectin-family, L-, P- and E-selectin, participate in multiple steps of the inflammatory cascade including the initial recruitment or tethering process, cell rolling, and the transition of rolling cells to firm arrest^12^. Here, P- and E-selectin expressed on the inflamed endothelium facilitate the direct recruitment of leukocytes from flowing blood, while L-selectin amplifies primary recruitment by enabling secondary leukocyte-leukocyte capture via L-selectin on one neutrophil engaging PSGL-1 (P-selectin glycoprotein ligand-1, CD162) on a second immune cell. Neutrophil capture and rolling is followed by chemokine driven cell activation, and firm arrest via members of the CD18/β_2_-integrin family (LFA-1, Mac-1)^13^.

Whereas the availability of blocking monoclonal antibodies has demonstrated that PSGL-1 is a major L- and P-selectin ligand on human neutrophils^14^, equivalent E-selectin ligands remain ill-defined^15^. In general, specific glycoforms of CD43, CD44 and CD162 are considered to be prime candidates for binding E-selectin on human immune cells^16^, as similar ligands have been identified in mice^17^. These glycoproteins bear O- and N-linked glycans containing sialofucosylated glycans like sialyl Lewis-X (Neu5Acα(2-3)Gal(β1-4)[Fucα1-3)]GlcNAcβ, CD15s) that engage the selectins efficiently under physiological fluid shear. While established in mice, it is recognized that mouse-human cross-species differences exist with respect to glycosylation pathways and thus findings from one species cannot be directly translated to another^15^. In this regard, a major mouse E-selectin ligand called ESL-1 (E-selectin ligand-1) has no equivalent in humans, and glycolipids enable slow rolling of human but not mouse neutrophils^18,19^. Additionally, distinct E-selectin ligands have been proposed to exist in different human immune cell populations^20^, though these propositions await validation using loss-of-function studies as specific blocking antibodies or genome editing strategies have not been applied. These technological problems are only exacerbated in peripheral blood neutrophils due to their short half-life, and thus the identity of E-selectin ligands on human neutrophils remains to be determined.

The short life and easy activation of human neutrophils in *ex vivo* culture represents a challenge that limits the study of these cells in their native state, especially under physiologically relevant hydrodynamic/fluid shear conditions. In this study, we address the possibility that such studies can be enabled by starting with primary CD34+ human hematopoietic stem/progenitor cells (‘hHSC’) and differentiating them to neutrophil-like cells, while simultaneously enabling CRISPR-Cas9 based gene editing. Using this approach, we established a protocol for studying E-selectin ligands on human neutrophils. Our findings suggest that a sialofucosylated glycoform of CD44 is a major E-selectin ligand on human neutrophils.

## MATERIALS AND METHODS

### Materials

Sources of commercial reagents are provided in Supplemental Methods. Wild-type HEK293T human embryonic kidney cells (‘HEK’) and HEK293Ts stably over-expressing the human α(1,3)fucosyltrasferase FUT7 as a FUT7-DsRed fusion protein (‘HEK-FUT7’) were available from previous studies^21,22^.

### Primary human blood neutrophils

Blood was collected from healthy human volunteers using venipuncture into 10U/mL heparin following protocols approved by the University at Buffalo Health Sciences Institutional Review Board (UB-HSIRB). Neutrophils were isolated using a one-step centrifugal process using the lympholyte-poly gradient buffer (Cedarlane, Burlington, ON)^23^. In some cases, these cells were cultured in complete IMDM medium with 10% fetal bovine serum, 10ng/mL recombinant human Interleukin-3 (IL-3) and 10ng/mL recombinant human Granulocyte Colony-Stimulating Factor (GCSF) for 2-4 days.

### CD34+ human hematopoietic cell isolation and culture

Deidentified, discarded human umbilical cord and placental blood was obtained from the Department of Obstetrics & Gynecology (Millard Fillmore Suburban Hospital, Williamsville, NY). CD34+ human hematopoietic stem/progenitor cells (hHCs) were isolated using the EasySep™ Human CD34 positive selection kit according to manufacturer’s instructions (StemCell Technology, Vancouver, Canada). These cells were cultured *ex vivo* using various protocols. In the ‘Base’ protocol, CD34+cells were cultured for 2 days in IMDM growth medium with 50ng/mL recombinant human Stem Cell Factor (SCF) and 50ng/mL IL-3. The amount of SCF and IL-3 was then reduced by 50% every 2 days thereafter. SCF and IL-3 levels were maintained at 10ng/mL until day10, after which it was removed. 10ng/mL GCSF was added to the growth medium between days 7-10, after which it was increased to 25ng/mL until day 14. Three modifications of the ‘Base’ protocol were also implemented: i. ‘+GM-CSF’: In addition to ‘Base’, 10ng/mL human granulocyte-macrophage colony-stimulating factor (GM-CSF) was added to the culture medium from day 7-10, followed by an increase to 25ng/mL from day 10-14. ii. ‘+TPO’: The ‘Base’ protocol was supplemented with 10ng/mL recombinant thrombopoietin (TPO) from day 0-7. iii. ‘+Flt-3L’: FMS-like tyrosine kinase 3 ligand (FLT3L) was added to the ‘Base at 10ng/mL prior to day 7. The differentiated cells were analyzed using cytospin analysis, RT-PCR and flow cytometry (Supplemental Methods).

### CRISPR-Cas9 genome editing of HL-60 cells and CD34+ hHSC derived neutrophils

sgRNA used for genome editing were either in native form or chemically modified to increase stability (**Supplemental Table S2**). In some cases, sgRNA was produced using *in vitro* transcription (Supplemental Methods), and in other cases chemically modified sgRNA were purchased from Synthego (Silicon Valley, CA). Target guide sequences were from published literature ^24,25^.

To knockout genes in leukocytes, ribonucleoprotein particles (RNP) were first formed by mixing 1μg sgRNA/gene with 1μg Cas9 (Cas9 NLS, *S. pyogenes*, New England Biolabs) at room temperature for 25-30 min. This was then added to 0.15-0.25×10^6^ cells suspended in 10μl Resuspension Buffer T provided with the Neon transfection kit (Thermo). Following this, the cell mixture was electroporated under optimized condition for all cell types: 1600V, 10ms, 3 pulses, and then transferred into 24-well plates containing 500μl of conditioned IMDM culture medium.

### CD44-Fc expression and immobilization on polystyrene beads

**‘**CD44-isoform1-CD4d3+4-bio’ plasmid was obtained from Addgene (# 73098, Watertown, MA^26^). The extracellular region of CD44 was PCR amplified from this construct using primers listed in **Supplementary Table S3**. This was ligated into a pCSCG plasmid available from a previous study, just upstream of the human Fc region^27^. This plasmid was transiently transfected into HEK and HEK-FUT7 cells using the calcium-phosphate method in the presence of serum-free medium^28^. Cell culture supernatant was harvested 72h post-transfection. Protein production was confirmed using western blotting and flow cytometry (Supplemental Methods).

### Cell adhesion assay

Primary neutrophils isolated from peripheral blood, neutrophils differentiated from CD34+ hHSCs, and beads coupled with CD44-Fc were washed and re-suspended in HEPES buffer containing 1.5mM CaCl2 and 0.1% human serum albumin (HSA). Plates were coated overnight at 4⁰C with either 1μg/ml recombinant E-selectin-IgG, 2μg/ml P-selectin-IgG or 10μg/ml L-selectin-IgG, and the substrate was blocked using PBS containing 3% BSA for 1h at RT. In some cases, HUVECs (human umbilical vein endothelial cells) were cultured in 35 x 10 mm tissue culture dishes until reaching 90∼100% confluence. The cell monolayer was then stimulated with 6.25U/mL IL-1β (R&D systems, Minneapolis, MN) for 4h at RT to upregulate E-selectin. 2×10^6^ cells/ml or 5×10^6^ beads/ml, suspended in HEPES+1.5mM Ca^2+^+0.1% HSA, were perfused at 0.5-1 dyne/cm^2^ over the selectin bearing substrate. Data were acquired at 2 frames/sec for 4min, using an inverted microscope. Cell rolling, firm adhesion density and cell rolling velocity were quantified as described previously^18^. When necessary, 10μg/ml receptor blocking mAbs were added to the substrate for 20 min, prior to cell/bead perfusion. For the CD44-FUT7-Fc blocking study, CD44-FUT7-Fc collected from cell culture medium was concentrated 200-fold using Amicon® Ultra-15 centrifugal filter units (Millipore Sigma), and 10µl of this concentrate was added to block the HUVEC substrate for 1h before performing the rolling assay at 1dyn/cm^2^. 10× CD44-FUT7-Fc (i.e 1:20 of the 200X concentrate) was also added to the flowing leukocytes to compensate for any loss of CD44-FUT7-Fc that was flushed away during the flow assay.

In studies that evaluated the rolling phenotype of RNP edited neutrophils derived from CD34+ hHSCs, the microfluidics flow cell was mounted on the stage of a Zeiss LSM 710 confocal microscope (Plan-Apochromat 20x/0.8 M27 objectives, QUASAR PMT detector) to distinguish between the different cell types. Here, mAbs against CD43 (MEM59-FITC) and CD44 (BJ18-APC) were added to monitor the knockout status of individual interacting cells. Data were acquired at 1 frame/s. The number of rolling and adherent cells that expressed CD43 only (green), CD44 only (red), both receptors (yellow) and dual KOs (white) were counted by frame-by-frame analysis. Normalized cell interaction density was calculated to account for gene editing efficiency based on the equation: Normalized interacting cell density=Interacting cell density for specific cell type ÷ % of such cells in the flow input based on flow cytometry analysis. This was calculated for wild-type, single- and double-KOs.

### Statistics

Data are presented as mean ± standard deviation for N≥3. Dual treatment comparisons were performed using a two-tailed Student’s T-test. Multiple comparisons were performed using ANOVA followed by the Tukey post-test. *P*<0.05 was considered to be significant.

### Data sharing

All data are provided in the main manuscript and data supplement available with the online version of this article.

## RESULTS

### Genome editing is not feasible using neutrophils isolated from peripheral blood

To study selectin-ligands on human neutrophils, previous studies cultured these cells *ex vivo* for 48h while performing chemical treatments, prior to functional testing^19^. We determined if this approach can be extended for genome engineering of primary neutrophils. Thus, human neutrophils isolated by gradient centrifugation from peripheral blood were cultured for 2-days in media containing IL-3 and G-CSF. Here, we observed a substantial loss of proliferating cells within 2-days (**Fig. 1A**). A majority of these cells, at day 2, also displayed low LDS-751 staining and high Annexin-V binding indicating increased cellular apoptosis (**Fig. 1A**). Consistent with this, no viable cells were detected on day 4.

**Figure 1.**
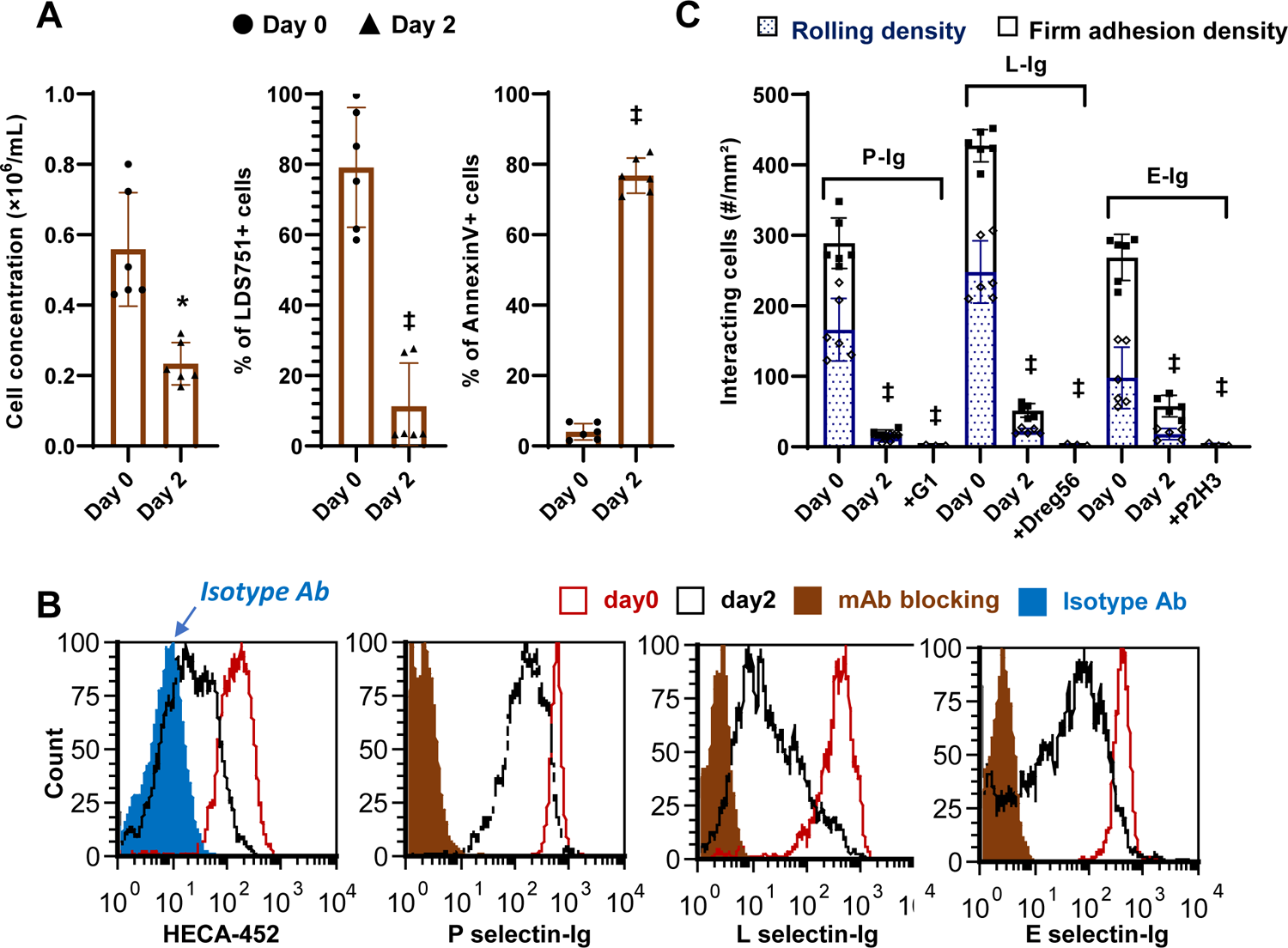
*Ex vivo* cultured peripheral blood neutrophil binding to selectins. **A.** Peripheral blood neutrophils were cultured in complete IMDM with 10 ng/mL IL-3 and 10ng/mL G-CSF. Hemocytometer based cell count decreased upon culture. Live cell staining using LDS-751 is decreased and Annexin-V positive apoptotic cell counts increased in 2 days. No viable cells remained at day 4. **B.** HECA-452 epitope expression decreased over time in culture. Binding to recombinant selectin-IgG proteins in flow cytometry assays was also reduced. **C.** Neutrophil rolling and firm adhesion in microfluidic flow assay was reduced upon culture for all selectins. Blocking mAbs were added to day 0 samples in panels B and C. These include clones G1 (anti-P-selectin), P2H3 (anti-E-selectin) and DREG-56 (anti-L-selectin), respectively. Data are mean ± S.D. for three independent donors (2-3 repeats/donor). * P<0.05, ‡ P<0.001 with respect to day 0.

Besides increased apoptosis, *ex vivo* culture of blood neutrophils also resulted in severe loss of selectin binding function. In this regard, a ∼70% reduction in cell surface sialyl Lewis-X (sLe^X^) expression occurred at day 2 based on mAb HECA-452 binding (**Fig. 1B**). Human selectin-IgG1 fusion proteins binding to cultured neutrophils was also reduced by 75% for P-selectin, 80% for L-selectin and 60% for E-selectin-IgG (**Fig. 1B**). Finally, there was dramatic loss of neutrophil binding function to substrates bearing all three selectins in cell adhesion assays performed under fluid shear (**Fig.1C**). Thus, alternate techniques are needed to perform genome engineering of primary human neutrophils.

### Obtaining human neutrophils upon differentiating CD34+ human hematopoietic cells

Genome editing requires several days of cell culture due to the time required to turnover and renew cell surface glycoproteins. We tested the possibility that such manipulations can be performed using neutrophils derived from CD34+ hHSCs. To this end, CD34+ hHSCs were isolated from human cord and placental blood, and then differentiated toward neutrophils by varying the cytokine cocktail added to culture medium (**Fig. 2A**). Four differentiation protocols were tested based on prior work that demonstrates the role of six chemokines during *ex vivo* expansion of hHSCs^29–32^. This includes the ‘Base’ protocol, where SCF and IL-3 were gradually decreased in the first 10 days prior to raising G-CSF concentration to drive differentiation to terminal neutrophils. This ‘Base’ method was additionally supplemented with either GM-CSF at later times (‘+ GMCSF’), TPO (‘+ TPO’) in the first 7 days or Flt-3L (+ ‘Flt3-L’) in the first week (**Fig. 2A**).

**Figure 2.**
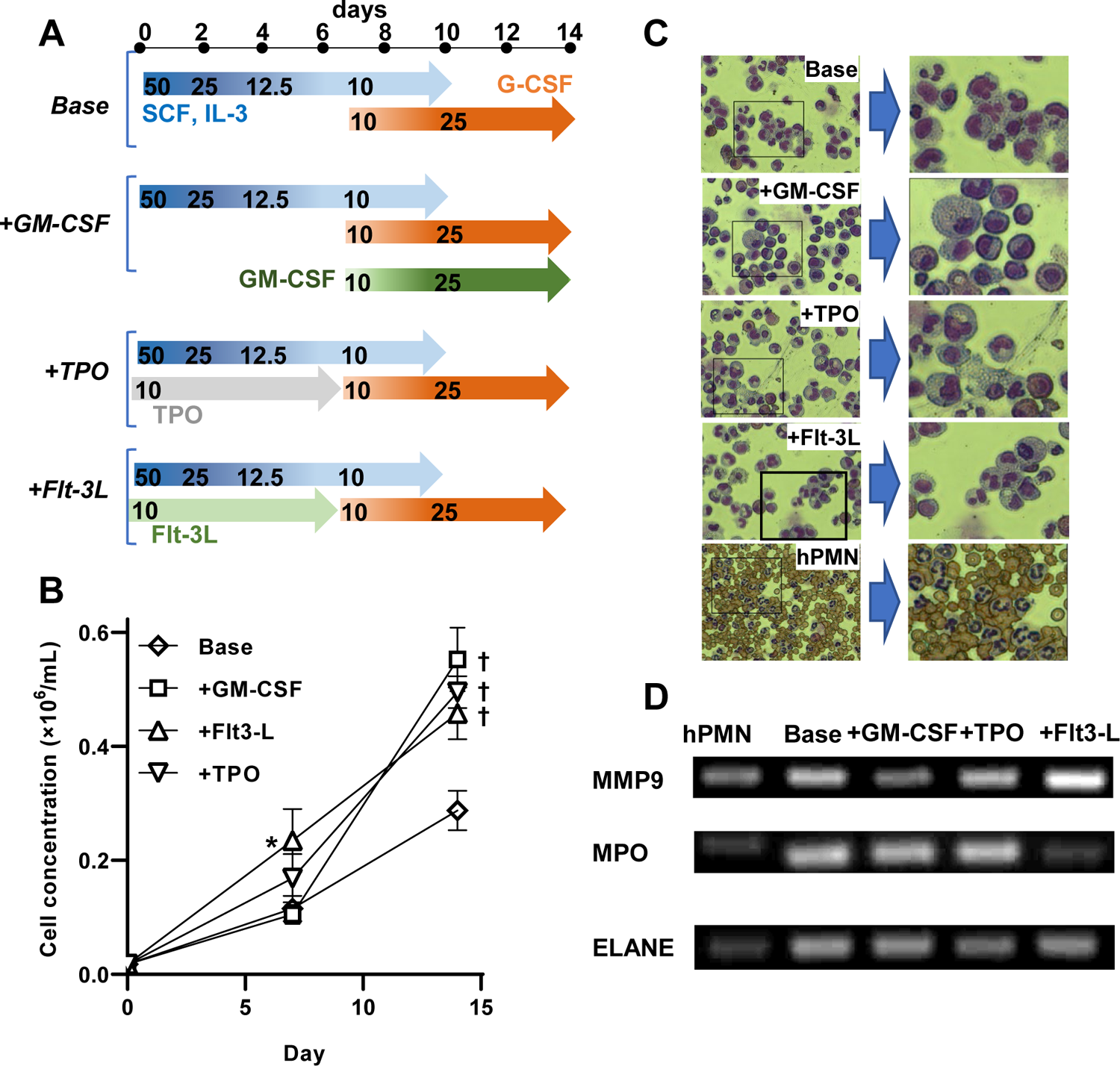
CD34+ human hematopoietic stem cell (hHSC) differentiation to primary neutrophils. **A.** Four differentiation protocols were tested to obtain mature neutrophils. The ‘Base’ includes graded application of SCF, IL-3 and G-CSF. Other protocols included ‘Base’ along with supplemental addition of either GM-CSF, TPO or Flt-3L. **B.** Cell proliferation data demonstrate increased cell numbers upon supplementation of the ‘Base’ cocktail with additional chemokines. **C.** Cytospins at day14 enumerate the number of cells with multi-lobed nucleus following differentiation. These varied as +Flt3-L (44%) > Base (30%) > +TPO (25%) > +GM-CSF (6%). **D.** RT-PCR amplicons compare expression levels of MPO (myeloperoxidase), MMP9 and ELANE (elastase) across the differentiation protocols. Data are mean ± S.D. for three independent donors (2-3 repeats/donor). * P<0.05, † P<0.01 with respect to ‘Base’ treatment. Cytospin and RT-PCR data are representative of >6 repeats.

The ‘Base’ protocol resulted in ∼30-fold expansion of hHSCs over 2-weeks, with additional cytokine supplementation resulting in double the number of output cells (**Fig. 2B**). Thus, upon starting with 100-170×10^3^ CD34+ HSCs on day 0, the final yield was typically ∼8-10×10^6^ cells. Cytospin analysis on day 14 enumerated cells with multi-lobed nuclei, which identify terminally-differentiated neutrophils, pre-mature band cells and related myelocyte states^33^. Here, ‘+Flt3-L’ resulted in the highest percentage of multi-lobed nucleated cells (44%), followed by ‘Base’ (30%), ‘+TPO’ (25%) and ‘+GM-CSF’ (6%) (**Fig. 2C**). While continued cell culture beyond day 14 resulted in marginal increase in cells expressing multi-lobed nucleus, this also resulted in loss of viability. The final cells obtained using these methods were larger (17-20 µm) compared to peripheral blood neutrophils (12-14 µm). Regardless of this size difference, the differentiated cells expressed a similar profile of several neutrophil genes based on RT-PCR including Matrix metallopeptidase 9 (MMP9, tertiary granule marker), Neutrophil elastase (ELANE, primary/azurophilic granules), and Myeloperoxidase (MPO, primary/azurophilic granules). (**Fig. 2D**, **Supplemental Table S1**). Cell structure and RT-PCR profile were particularly similar for cells obtained using the ‘+Flt3-L’ protocol, compared to blood neutrophils.

Myeloid cell surface markers were measured using flow cytometry on day 7 and 14 to characterize the differentiation protocol (**Fig. 3**). Here, the measured markers were typically lower on day 7 compared to day 14, indicating cell maturation in the final stretch. CD15 levels at day 14 were comparable to that of neutrophils (**Fig. 3A**). CD11b (**Fig. 3B**) and CD33 (**Fig. 3C**) were expressed at higher level on the differentiated cells, and CD16 was lower (**Fig. 3D**). The HECA-452 epitope was low at day 7 but markedly increased at later times, consistent with prior reports indicating an increase in α(1,3)fucosylation in the last stages of neutrophil maturation^34^ (**Fig. 3E**). Most of the measured receptor expression levels were uniform across the entire differentiated cell population, though two CD11b populations were detected at day 14 (**Supplemental Fig. S1**). Overall, a protocol for *ex vivo* hHSC differentiation was optimized to yield ∼50-80 fold CD34+ hHSC expansion over a 2-week period. The derived neutrophils expressed the expected cell surface markers with morphological features resembling terminally differentiated neutrophils.

**Figure 3.**
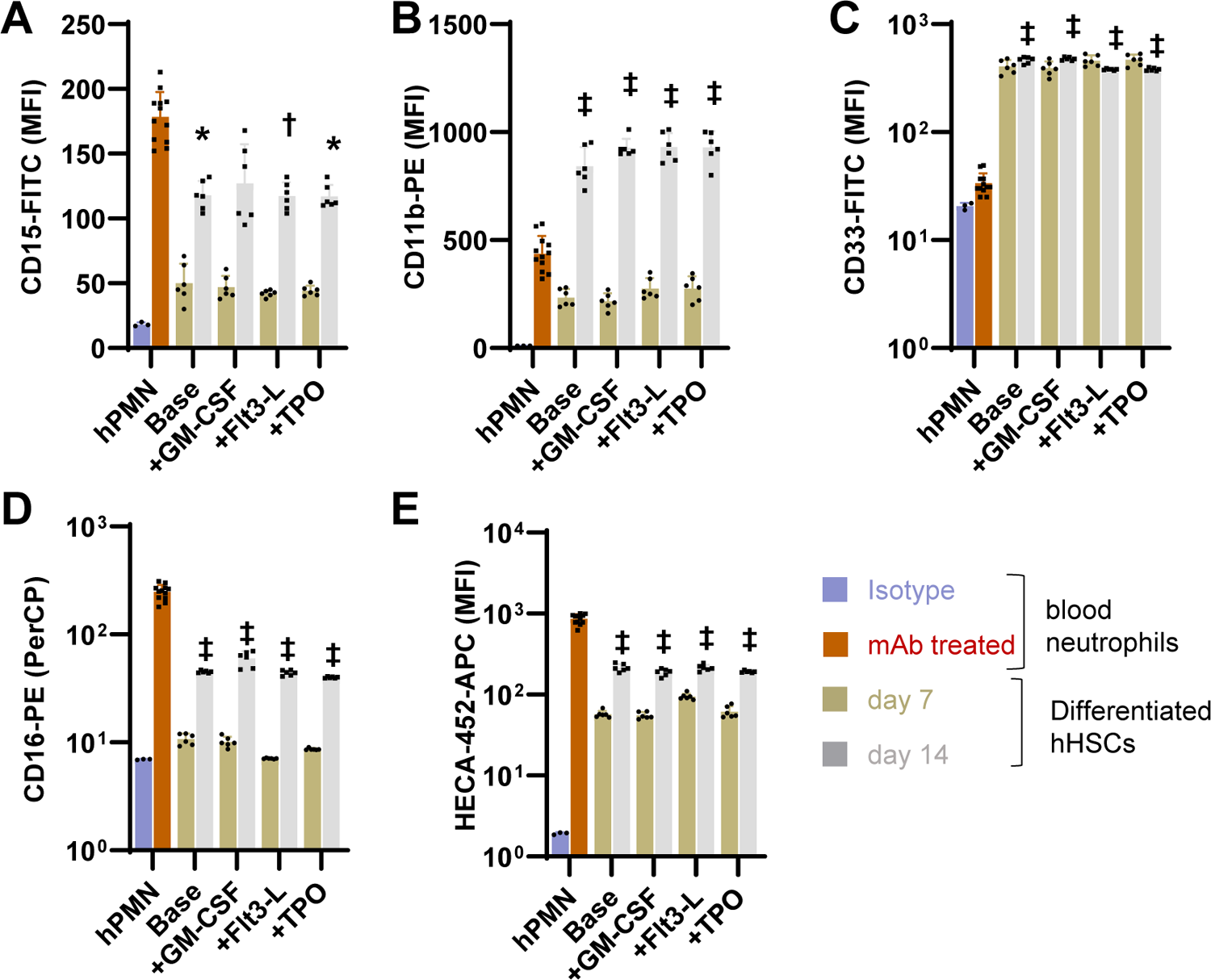
Epitope expression on neutrophils differentiated from hHSCs. Flow cytometry measured various myeloid markers in the differentiated cells, and compared them to peripheral blood neutrophils. Regardless of the differentiation protocol, neutrophils derived from hHSCs displayed CD15 levels comparable to primary neutrophils (panel A), higher CD11b (B) and CD33 (C) expression, and reduced CD16 levels. The sLe^X^/CLA epitope recognized by mAb HECA-452 increased markedly from day 7 to 14, and approached blood neutrophil levels at the latter time point. Thus, the upregulation of sLe^X^ expression is a late-stage differentiation event. MFI: Mean fluorescence intensity. Data are presented as mean ± S.D. for 3 independent donors (each repeated 4-6 times). Statistics for day 14 differentiated cells with respect to peripheral neutrophils: * P<0.05, † P<0.01, ‡ P<0.001.

### Neutrophils differentiated from hHSCs displayed robust selectin-ligand binding function

Selectin-dependent binding assays were performed to confirm glycan dependent cell adhesion function. In static assays, recombinant human selectin-IgG were complexed with a secondary fluorescent antibody, prior to addition to peripheral blood neutrophils or neutrophils differentiated from CD34+ hHSCs. Here, we observed robust binding of P-(**Fig. 4A**) and L-selectin IgG (**Fig. 4B**) to all cell types at early times (day 7), with the absolute levels of selectin-IgG binding being like that of blood neutrophils by day 14. The binding pattern was similar for both P- and L-selectin as they both primarily recognize neutrophil PSGL-1. In contrast, E-selectin IgG binding to differentiated cells was low at day 7 consistent with the low expression of the HECA-452 epitope (**Fig. 4C**, **Supplemental Fig. S2**). Both E-selectin-IgG1 and HECA-452 binding increased dramatically by day 14. The pattern of selectin-binding was not very different among the differentiation protocols. In controls, selectin binding was completely blocked by anti-selectin mAbs against P-selectin (clone G1), L-selectin (Dreg56) and E-selectin (P2H3).

**Figure 4.**
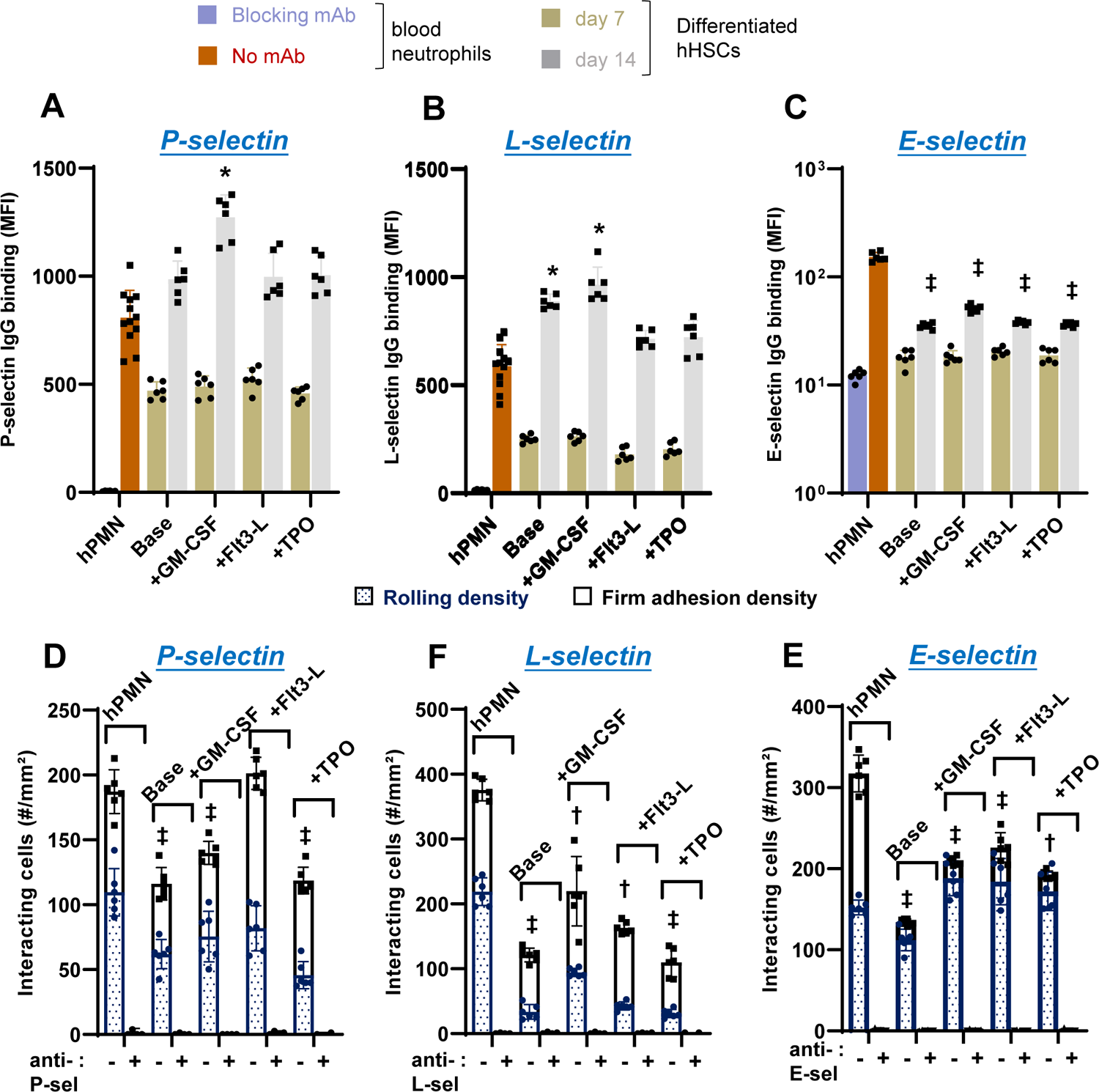
Neutrophil binding to selectins under static and shear condition. **A-C**. Flow cytometry compared the binding of various cell types to P-/L-/E-selectin-IgG fusion protein under static conditions. P- and L-selectin IgG binding at day 14 were comparable to that of primary neutrophils. E-selectin Ig binding was lower, but increased from day 7 to 14. **D**-**F**. 2×10^6^ differentiated cells (day 14) were perfused over substrates bearing P-/L-/E-selectin under fluid shear conditions (1 dyn/cm^2^). Cell rolling density was slightly lower for the differentiated cells compared to primary neutrophils for all treatments. Binding specificity for static and shear assays was confirmed using blocking mAbs against P-, E- and L-selectin IgG (mAbs G1, P2H3 and DREG-56, respectively). Data are mean ± SD from 4-6 repeats. * P<0.05, † P<0.01, ‡ P<0.001 with respect to peripheral blood neutrophils.

In microfluidics flow chamber assays performed at 1 dyn/cm^2^, we observed robust rolling of the differentiated cells on all three selectin substrates, though the absolute number of interacting cells was lower compared to blood neutrophils (**Fig. 4D**-**F**). This may be due to the larger size of the differentiated cells which experience higher hydrodynamic drag forces. Among the treatments, ‘+Flt3L’ provided highest interaction with P- and E-selectin. Cell rolling velocity of all the differentiated cells derived from hHSCs was similar to that of blood neutrophils on P-/L-selectin, with rolling velocity being two-times greater for the differentiated cells on substrates bearing E-selectin (**Supplemental Fig. S3**). In controls, cell adhesion were blocked using anti-selectin mAbs. Overall, neutrophils derived using the ‘+Flt3L’ protocol most closely resembled peripheral blood neutrophils in terms of the multi-lobed nuclear morphology, gene expression profile and rolling phenotype on E-/P-selectin. Thus, this protocol was adopted for all studies described below.

### Ribonucleoprotein complex based gene editing of differentiated neutrophils

Neutrophils derived from hHSCs were hard-to-transfect using conventional plasmid-based methods and cell viability was poor upon using lentivirus (data not shown). Thus, we adopted a previously described ribonucleoprotein particle (RNP) based method for gene editing^35^.

In one study, sgRNA were synthesized using *in vitro* transcription, and these were complexed with *S. pyrogenes* Cas9 protein (**Fig. 5A**). 5-10 sgRNA were tested for each target, and the final/most-efficient selection is listed in **Supplemental Table S2**. Electroporation of RNPs formed using these sgRNA resulted in >95% reduction in CD43 and CD44 expression in HL-60 cells. Remarkably, however, sgRNA editing efficient was drastically reduced in cells derived from CD34+ hHSCs. Thus, the editing efficiency of the CD43 RNPs was reduced from 96% to 71% upon use in neutrophils derived using the ‘+Flt-3L’ protocol, and from 96% to 20% in the case of the CD44 RNPs. This reduced efficacy could be rescued upon using sgRNA protected by 2’-O-methyl 3’ phosphorothioate inter-nucleotide linkages at the three terminal 5’ and 3’ bases of the sgRNA, suggesting RNA degradation in primary neutrophils^36,37^. Such modified sgRNA yielded 65-70% and 75-80% knockout (KO) of CD43 and CD44, respectively, in single gene knockouts using neutrophils derived from CD34+ hHSCs at day 14 (**Supplemental Fig. S4**).

**Figure 5.**
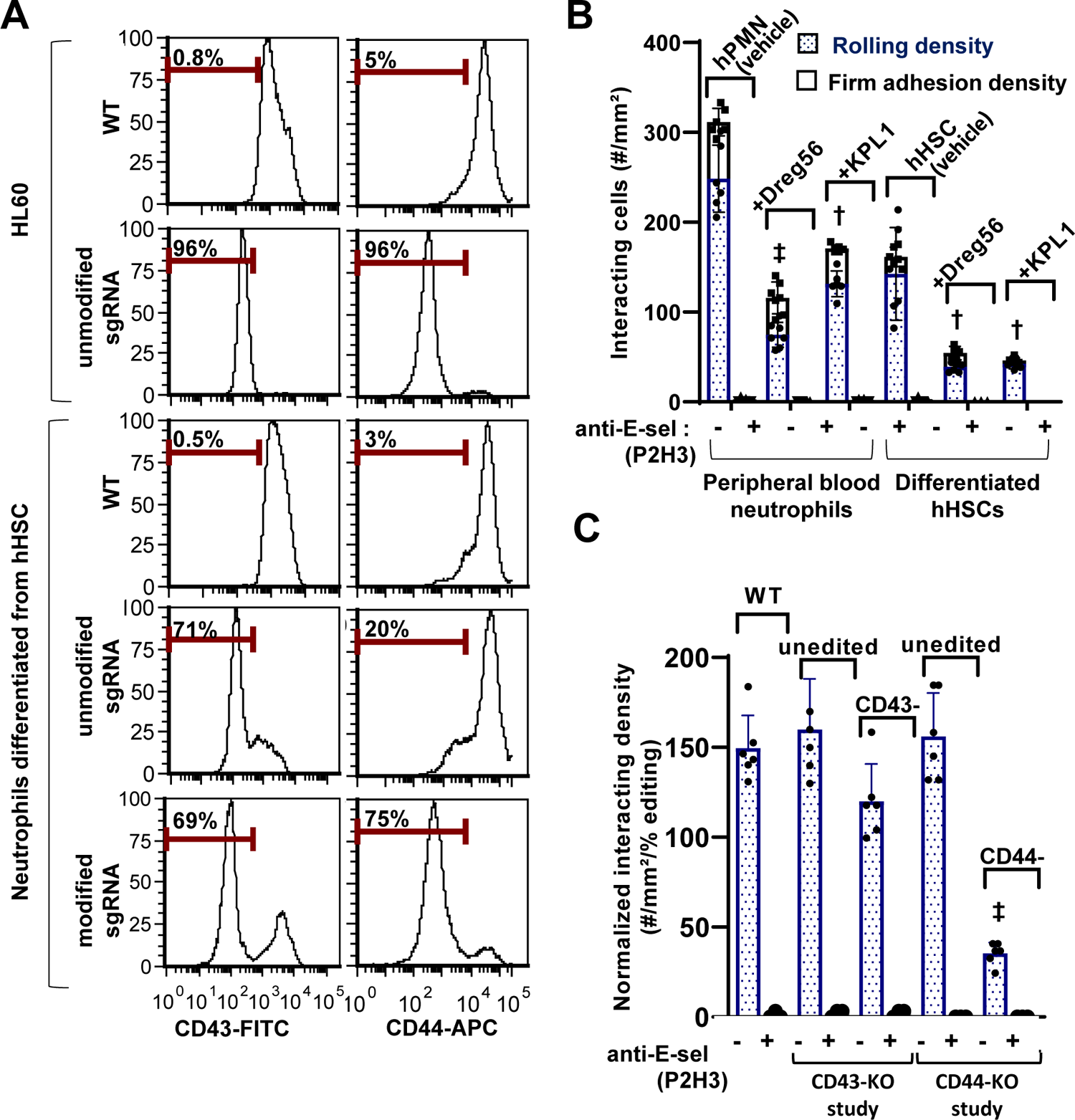
Gene editing of hHSC derived neutrophils using CRISPR-Cas9. **A.** Ribonucleoprotein (RNP) complexes were generated by mixing 1µg Cas9 and 1µg sgRNA (unmodified or modified) targeting either CD43 (left column) or CD44 (right column) in 2µL volume. RNPs formed were electroporation into 0.1-0.25×10^6^ HL60s (top) or neutrophils derived from hHSCs (bottom). Flow cytometry measured receptor expression levels at 48h. RNA degradation was severe in neutrophils derived from hHSCs, and this could be rescued by using modified sgRNA. ∼70% single gene editing was typically possible in the neutrophils derived from hHSCs. **B.** Peripheral blood neutrophils or differentiated hHSCs were perfused over E-selectin substrates under flow at 1 dyn/cm^2^. Neutrophil-neutrophil interactions were reduced by addition of either anti-L-selectin (Dreg56) or anti-PSGL-1 (KPL-1) mAb. Binding interaction was E-selectin specific. **C.** Unedited neutrophils, CD43- and CD44-knockouts derived from hHSCs were perfused over substrates bearing E-selectin at 1 dyn/cm^2^, in the presence of 10µg/mL anti-CD18 and anti-PSGL-1 (KPL1) blocking mAbs. Cells were labeled with anti-CD43 FITC and anti-CD44 APC to distinguish unedited vs. edited cells. Rolling data were normalized based on editing efficiency. Cell rolling was characterized for paired unedited and edited cells in each case. Targeting CD44 significantly reduced cell rolling. Data are presented as mean ± S.D. for 4-6 independent repeats for each treatment. † P<0.01, ‡ P<0.001 with respect to unedited WT cells.

Cell adhesion studies evaluated the effect of single gene knockout on substrates composed of E-selectin (**Fig. 5B**, **5C**). First, neutrophils derived from peripheral blood and CD34+ hHSCs were perfused over this substrate (**Fig. 5B**). Here, we observed clusters of neutrophils rolling in proximity due to hydrodynamic effects and secondary neutrophil-neutrophil interaction (**Supplemental Movie A**). This was likely due to presentation of both L-selectin and PSGL-1 by the neutrophils. Consistent with this, similar proportions of secondary adhesion events could be blocked for both blood neutrophils and neutrophils derived from hHSCs, upon using either anti-PGSL-1 or anti-L-selectin blocking mAbs (**Fig. 5B**). In order to reduce secondary recruitment and prevent firm arrest, all subsequent studies were performed in the presence of anti-PSGL-1 and anti-CD18 mAbs. Additionally, the microfluidics flow chambers were mounted on a confocal microscope to assay the phenotype of the rolling cells. This was achieved by labeling blood cells with non-function blocking mAbs against CD43 (MEM59-FITC) and CD44 (BJ18-APC) and using fluorescence to monitor the phenotype of rolling cells. In a representative example, RNP directed against CD43 resulted in 68% gene editing as measured using flow cytometry, and the proportion of rolling/interacting KO cells was similar (64%) (**Supplemental Fig. S4**). In contrast, editing 80% of the blood cells using CD44 RNP resulted in only 22% of the rolling cells being CD44-KO. This suggests that CD44, but not CD43, is a potential human neutrophil E-selectin ligand. To account for KO-efficiency, thus, rolling data were normalized with respect to % editing (**Fig. 5C**).

### Studies with dual knockouts confirm the role of CD44 as a physiological human E-selectin ligand

To evaluated if CD43 may act in synergy with CD44, the RNPs were mixed to produce dual knockout cells (**Fig. 6A**). Here, in a representative run, we obtained 42% CD44/CD43 dual KOs (DKOs), 8% CD44-SKO (single-KO) and 30% CD43-SKO, with the remaining 20% cells being unedited. The proportional of cells rolling in the confocal microscopy setup was quite different with few interacting CD44-SKO and CD43/CD44-SKO cells on E-selectin (**Fig. 6B**, **Supplemental Movie B**). A synergistic role for CD43 was not observed, though the prominent role for CD44 was reconfirmed (**Fig. 6C**).

**Figure 6.**
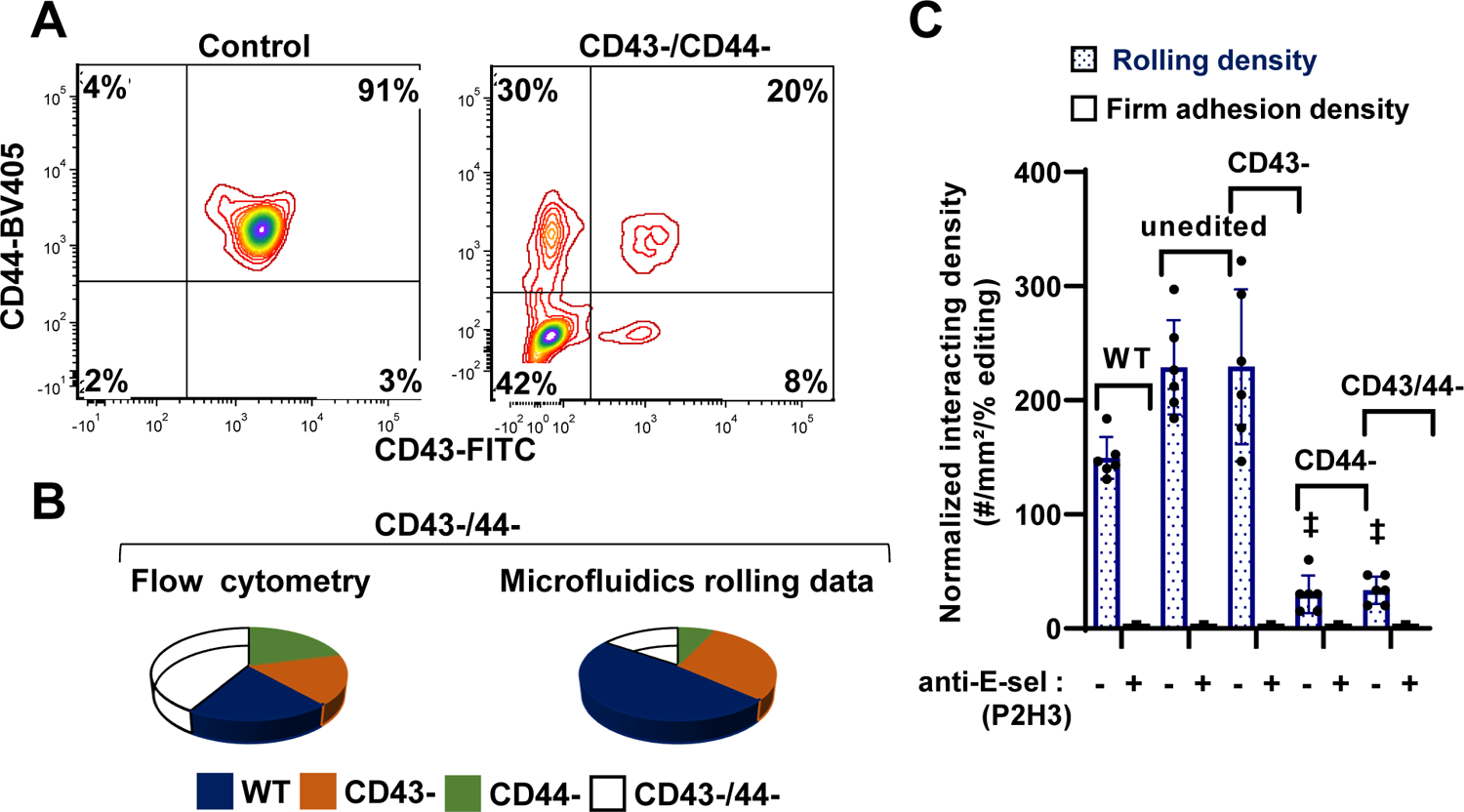
Confocal microfluidic rolling assay for mixture of multiple knockouts. **A**. RNPs targeting CD43 and CD44 were produced by mixing 1µg Cas9, 1µg CD43 modified sgRNA and 1µg CD44 modified sgRNA in 2µL volume. These were electroporated into hHSCs at day 2 followed by differentiation to neutrophils. Flow cytometry measured receptor expression at day 14 for cells electroporated with Cas9 alone (‘control’) and those containing electroporated with RNP mixture (‘CD43-/CD44-’). **B-C.** Cells labelled with anti CD43-FITC and anti CD44-APC were perfused over E-selectin substrates at 1 dyn/cm^2^. Cell rolling density was measured using confocal microscopy with antibody labels being used to distinguish wild-type cells from individual knockouts. Data are presented in the form of pie chart in panel B, in order to enumerate the different edited populations observed using flow cytometry vs. those observed under flow/shear. Rolling cell density are normalized based on editing efficiency measured using cytometry (panel C). Results show that CD44 is an important neutrophil E-selectin ligand. Data are presented as mean ± S.D. for 4-6 independent repeats for each treatment. ‡ P<0.001 with respect to unedited WT cells.

### Cell free assays reveal a crucial role for CD44 α(1,3)fucosylation and HECA-452 epitope for E-selectin mediated cell adhesion

Cell free rolling assays were performed to determine if sialofucosylated glycans on CD44 can alone contribute to leukocyte rolling. Thus, we expressed CD44 as a fusion protein with C-terminal human IgG1, in both wild-type HEK cells and also HEK cells stably over-expresses the human α(1,3)fucosyltransferase FUT7^22^. These proteins products are called ‘CD44-Fc’ and ‘CD44-FUT7-Fc’, respectively (**Fig. 7A**). Both proteins had molecular mass of ∼250kDa, with CD44-FUT7-Fc also expressing the sialyl Lewis-X epitope as recognized by mAb HECA-452 (**Fig. 7B**). Next, these proteins were immobilized on 6μm polystyrene beads that bear goat anti-human Ab at densities comparable to that of peripheral blood neutrophils (**Fig. 7C**, **Supplemental Fig. S5**). Among these constructs, beads bearing CD44-FUT7-Fc, but not CD44-Fc, displayed rolling phenotype on substrates bearing recombinant E-selectin (**Fig. 7D**, **Supplemental Movie C**). This specific interaction could be completely blocked by anti-human E-selectin mAb.

**Figure 7.**
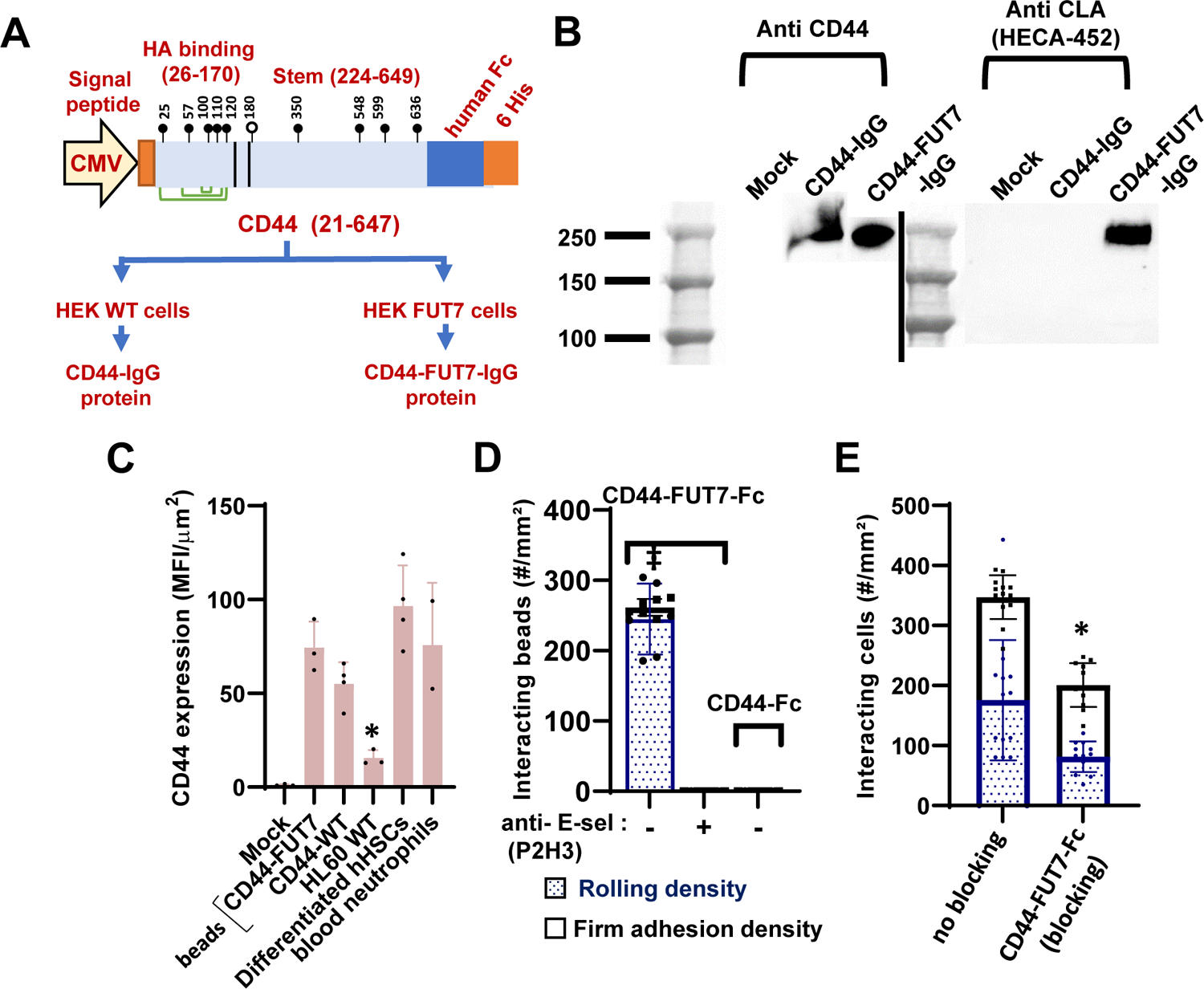
Cell free microfluidics rolling assay using α(1,3)fucosylated CD44-IgG. **A**. CD44-IgG-his tag protein was expressed in wild-type HEK293T cells to generated ‘CD44-IgG’ and HEK293T cells overexpressing the human α(1,3)fucosyltransferase FUT7 to produce ‘CD44-FUT7-IgG’. Open and closed lollipops indicate putative sites of N- and O-glycosylation. Green lines show disulfide bonds. **B.** Western blot of protein secreted from these cells, along with expression of HECA-452/sLe^X^ on these constructs. **C.** CD44-IgG and CD44-FUT7-IgG were immobilized on 5µm goat anti-human Ab bearing polystyrene beads. Immobilized CD44 concentration was measured using fluorescent anti-CD44 Ab. CD44 density on cells and beads are comparable after normalizing based on cell/bead size: differentiated hHSCs (18µm), blood neutrophils (13µm) and HL-60 leukocytes (17µm). CD44 density was lower on HL-60s, but comparable for other systems. **D.** CD44-FUT7-IgG but not CD44-IgG immobilized beads displayed robust rolling on substrates bearing recombinant E-selectin. This interaction could be blocked by anti-E-selectin mAb P2H3. **E.** Interaction between peripheral blood neutrophils and stimulated HUVEC cells was reduced by blocking the substrate bearing E-selectin with CD44-FUT7-Fc fusion protein. Wall shear stress=1 dyn/cm^2^ in all runs. Data are presented as mean ± S.D. typically for 4-6 independent repeats for each treatment. * P<0.05, ‡ P<0.001 with respect to all other treatments.

Next, human peripheral blood neutrophils isolated from healthy volunteers were perfused over substrates bearing stimulated HUVECs that express E-selectin (**Fig. 7E**). This cell rolling interaction was significantly blocked by CD44-FUT7-Fc but not vehicle control. Functional blocking in the context of vascular endothelial cells confirms the physiological importance of CD44 as a human neutrophil E-selectin ligand.

## DISCUSSION

Neutrophils play an essential role during acute and chronic inflammation with selectin-dependent adhesion being central to the initiation of cellular localization. This study established a methodology to perform gene editing on primary human neutrophil-like cells starting with CD34+ hHSCs. The final, optimized protocol using a cocktail of Flt-3L, IL-3, SCF and G-CSF resulted in 40% of the cells displaying multi-lobed nucleus. Among the cytokines used, SCF was used to promote hHSC proliferation and prevent apoptosis^38,39^, and IL-3 aided stem cell expansion^40^. Flt3-L^41–44^ along with G-CSF^45,46^ promoted colony formation and neutrophil differentiation. Other chemokines that we tested like TPO and GM-CSF reduced the formation of multi-lobed nucleated cells. Although the gene and protein expression profile of the final differentiated cells resembled blood neutrophils, detailed single-cell analysis in the future is needed for nuanced analysis of the cellular transcriptome. This was not pursued, due to greater interest in advancing gene editing methods and characterization of leukocyte adhesion mechanics. Our efforts resulted in a streamline protocol to study selectin-dependent cell adhesion, following >80% CRISPR-Cas9 based gene editing in hard-to-transfect, short-lived primary neutrophils.

A multicolor rolling assay was established on a confocal microscope to track the cell adhesion features of individual cell types in mixed populations. This approach demonstrated that a sialofucosylated glycoform of CD44 on human neutrophils is a major E-selectin ligand. Here, knocking out CD44 on primary neutrophils resulted in 75-80% reduction in cell adhesion on E-selectin under shear. Knocking out CD43, alone or in tandem with CD44, did not have a measurable effect suggesting that the CD43 glycoform on human neutrophils is not a physiological E-selectin ligand. In this regard, though a glycoform of CD43 has been proposed to act as a functional ligand for human monocytes and CD4+ T-cells^20^, this does not carry over to human neutrophils either because of different glycosylation machinery in these cells or due to the nature study design, as our conclusions are based on gene editing and loss-of-function assays under physiological shear, while Silvia *et al.*^20^ focus on expression levels of sialyl-Lewis X or E-selectin-IgG binding in immunoblots. In addition, we do not draw conclusions on the physiological role of human PSGL-1 E-selectin binding to neutrophil capture and rolling, as PSGL-1 predominantly contributed to secondary neutrophil-neutrophil capture. The same is true for L-selectin which has been proposed to bind E-selectin^47^. Finally, some residual leukocyte rolling (∼10-20%) was observed even upon blocking CD43 and CD44 using CRISPR-Cas9 and PSGL-1 using mAb, suggesting the presence of additional yet undetermined E-selectin ligands.

Functional selectin-like rolling could be reconstituted in cell-free assays under flow. α(1-3) fucosylation of CD44 was critical for such binding as only CD44 produced in HEK-FUT7 cells but not wild-type HEK cells were functional. In addition, sialofucosylated CD44-FUT7-Fc reduced neutrophil adhesion by 40-50% on E-selectin expressed on inflamed endothelial cells suggesting that this is physiologically relevant. In previous work, it has been proposed that both the N-glycans and O-glycans of CD44 may bind E-selectin, with the N-glycans being necessary for CD4+ T-cells and O-glycans being important for blood monocytes^20^. Cell-specific N- and O-glycans on individual CD44 isoforms of breast^48^ and colon^49^ cancer cells, have also been shown to bind endothelial E-selectin to facilitate metastasis. More detailed investigations are necessary to establish the exact contribution of these N-/O-glycoforms to human neutrophil function, and also to determine the precise glycosylation site that contributes to cell adhesion. Such efforts will necessitate the development of new glycoform-specific anti-CD44 mAbs that will reduce neutrophil binding to E-selectin under shear, as at least two anti-CD44 mAbs we tested (IM7 and Hermes-1) did not exhibit function-blocking properties in our assays. The development of such glycoform-specific mAbs could aid future CD44/E-selectin directed therapy for human inflammatory conditions.

Using HL-60 human promyelocytic cells as a model system, we previously noted a major role for N-glycans in facilitating myeloid cell capture and rolling on E-selectin, with leukocyte O-glycans also facilitating the initial tethering^12^. Using shRNA in HL-60s and also human neutrophils, we also established a functional role for glycolipids in the transition of neutrophils to firm arrest on the vascular endothelium^18^. Based on the current study, our working hypothesis is that O-glycans on PSGL-1 and N-glycans on CD44 may facilitate human neutrophil capture on endothelial cells, with the CD44 N-glycans also coordinating stable rolling and the transition to glycosphingolipid mediated slow rolling. The tools established in this paper will be used to test this model. Overall, this paper establishes CRISPR-Cas9 methods for studies of neutrophil biology that could be broadly useful beyond the investigation of selectins.

## ACKNOWLEDGEMENT

This work was supported by the National Heart, Lung and Blood Institute (NHLBI) Systems Biology grant HL103411 (S.N.).

## AUTHORSHIP

Y.Z.: Designed study, performed experiments, analyzed data and prepared original draft of manuscript. S.N: Designed study, supervised project and prepared final manuscript.

## CONFLICT OF INTEREST STATEMENT

The authors declare no competing financial interests.

## Notes

### Competing Interest Statement

The authors have declared no competing interest.

## REFERENCES

1. Lahoz-Beneytez J, Elemans M, Zhang Y, et al. Human neutrophil kinetics: modeling of stable isotope labeling data supports short blood neutrophil half-lives. Blood. 2016;127(26):3431–3438.

2. Herrero-Cervera A, Soehnlein O, Kenne E. Neutrophils in chronic inflammatory diseases. Cell Mol Immunol. 2022;19(2):177–191.

3. Bert S, Nadkarni S, Perretti M. Neutrophil-T cell crosstalk and the control of the host inflammatory response. Immunol Rev. 2023;314(1):36–49.

4. Silvestre-Roig C, Fridlender ZG, Glogauer M, Scapini P. Neutrophil Diversity in Health and Disease. Trends Immunol. 2019;40(7):565–583.

5. Liew PX, Kubes P. The Neutrophil’s Role During Health and Disease. Physiol Rev. 2019;99(2):1223–1248.

6. Grecian R, Whyte MKB, Walmsley SR. The role of neutrophils in cancer. Br Med Bull. 2018;128(1):5–14.

7. Gao D, Mittal V. The role of bone-marrow-derived cells in tumor growth, metastasis initiation and progression. Trends Mol Med. 2009;15(8):333–343.

8. Murdoch C, Muthana M, Coffelt SB, Lewis CE. The role of myeloid cells in the promotion of tumour angiogenesis. Nat Rev Cancer. 2008;8(8):618–631.

9. Kitamura T, Qian BZ, Pollard JW. Immune cell promotion of metastasis. Nat Rev Immunol. 2015;15(2):73–86.

10. Engblom C, Pfirschke C, Pittet MJ. The role of myeloid cells in cancer therapies. Nat Rev Cancer. 2016;16(7):447–462.

11. Buffone A, Jr., Nasirikenari M, Manhardt CT, et al. Leukocyte-borne alpha(1,3)-fucose is a negative regulator of beta2-integrin-dependent recruitment in lung inflammation. J Leukoc Biol. 2017;101(2):459–470.

12. Stolfa G, Mondal N, Zhu Y, Yu X, Buffone A, Jr., Neelamegham S. Using CRISPR-Cas9 to quantify the contributions of O-glycans, N-glycans and Glycosphingolipids to human leukocyte-endothelium adhesion. Sci Rep. 2016;6:30392.

13. Nourshargh S, Alon R. Leukocyte migration into inflamed tissues. Immunity. 2014;41(5):694–707.

14. McEver RP, Cummings RD. Perspectives series: cell adhesion in vascular biology. Role of PSGL-1 binding to selectins in leukocyte recruitment. J Clin Invest. 1997;100(3):485–491.

15. Mondal N, Buffone A, Jr., Neelamegham S. Distinct glycosyltransferases synthesize E-selectin ligands in human vs. mouse leukocytes. Cell Adh Migr. 2013;7(3):288–292.

16. Merzaban JS, Burdick MM, Gadhoum SZ, et al. Analysis of glycoprotein E-selectin ligands on human and mouse marrow cells enriched for hematopoietic stem/progenitor cells. Blood. 2011;118(7):1774–1783.

17. Hidalgo A, Peired AJ, Wild M, Vestweber D, Frenette PS. Complete identification of E-selectin ligands on neutrophils reveals distinct functions of PSGL-1, ESL-1, and CD44. Immunity. 2007;26(4):477–489.

18. Mondal N, Stolfa G, Antonopoulos A, et al. Glycosphingolipids on Human Myeloid Cells Stabilize E-Selectin-Dependent Rolling in the Multistep Leukocyte Adhesion Cascade. Arterioscler Thromb Vasc Biol. 2016;36(4):718–727.

19. Nimrichter L, Burdick MM, Aoki K, et al. E-selectin receptors on human leukocytes. Blood. 2008;112(9):3744–3752.

20. Silva M, Fung RKF, Donnelly CB, Videira PA, Sackstein R. Cell-Specific Variation in E-Selectin Ligand Expression among Human Peripheral Blood Mononuclear Cells: Implications for Immunosurveillance and Pathobiology. J Immunol. 2017;198(9):3576–3587.

21. Lo CY, Antonopoulos A, Gupta R, et al. Competition between core-2 GlcNAc-transferase and ST6GalNAc-transferase regulates the synthesis of the leukocyte selectin ligand on human P-selectin glycoprotein ligand-1. J Biol Chem. 2013;288(20):13974–13987.

22. Lo CY, Antonopoulos A, Dell A, Haslam SM, Lee T, Neelamegham S. The use of surface immobilization of P-selectin glycoprotein ligand-1 on mesenchymal stem cells to facilitate selectin mediated cell tethering and rolling. Biomaterials. 2013;34(33):8213–8222.

23. Zhang Y, Neelamegham S. Estimating the efficiency of cell capture and arrest in flow chambers: study of neutrophil binding via E-selectin and ICAM-1. Biophys J. 2002;83(4):1934–1952.

24. Tzelepis K, Koike-Yusa H, De Braekeleer E, et al. A CRISPR Dropout Screen Identifies Genetic Vulnerabilities and Therapeutic Targets in Acute Myeloid Leukemia. Cell Rep. 2016;17(4):1193–1205.

25. Wang T, Birsoy K, Hughes NW, et al. Identification and characterization of essential genes in the human genome. Science. 2015;350(6264):1096-1101.

26. Crosnier C, Bustamante LY, Bartholdson SJ, et al. Basigin is a receptor essential for erythrocyte invasion by Plasmodium falciparum. Nature. 2011;480(7378):534-U158.

27. Lo CY, Antonopoulos A, Dell A, Haslam SM, Lee T, Neelamegham S. The use of surface immobilization of P-selectin glycoprotein ligand-1 on mesenchymal stem cells to facilitate selectin mediated cell tethering and rolling. Biomaterials. 2013;34(33):8213–8222.

28. Buffone A, Jr., Mondal N, Gupta R, McHugh KP, Lau JT, Neelamegham S. Silencing alpha1,3-fucosyltransferases in human leukocytes reveals a role for FUT9 enzyme during E-selectin-mediated cell adhesion. J Biol Chem. 2013;288(3):1620–1633.

29. Sauvageau G, Iscove NN, Humphries RK. In vitro and in vivo expansion of hematopoietic stem cells. Oncogene. 2004;23(43):7223–7232.

30. Metcalf D. Hematopoietic cytokines. Blood. 2008;111(2):485–491.

31. Tajer P, Pike-Overzet K, Arias S, Havenga M, Staal FJT. Ex Vivo Expansion of Hematopoietic Stem Cells for Therapeutic Purposes: Lessons from Development and the Niche. Cells. 2019;8(2).

32. Ko KH, Nordon R, O’Brien TA, Symonds G, Dolnikov A. Ex Vivo Expansion of Hematopoietic Stem Cells to Improve Engraftment in Stem Cell Transplantation. Methods Mol Biol. 2017;1524:301–311.

33. Pillay J, Tak T, Kamp VM, Koenderman L. Immune suppression by neutrophils and granulocytic myeloid-derived suppressor cells: similarities and differences. Cell Mol Life Sci. 2013;70(20):3813–3827.

34. Mondal N, Dykstra B, Lee J, et al. Distinct human (1,3)-fucosyltransferases drive Lewis-X/sialyl Lewis-X assembly in human cells. Journal of Biological Chemistry. 2018;293(19):7300–7314.

35. Gundry MC, Brunetti L, Lin A, et al. Highly Efficient Genome Editing of Murine and Human Hematopoietic Progenitor Cells by CRISPR/Cas9. Cell Rep. 2016;17(5):1453–1461.

36. Hendel A, Bak RO, Clark JT, et al. Chemically modified guide RNAs enhance CRISPR-Cas genome editing in human primary cells. Nat Biotechnol. 2015;33(9):985–989.

37. Bak RO, Dever DP, Porteus MH. CRISPR/Cas9 genome editing in human hematopoietic stem cells. Nat Protoc. 2018;13(2):358–376.

38. Zhang CC, Lodish HF. Cytokines regulating hematopoietic stem cell function. Curr Opin Hematol. 2008;15(4):307–311.

39. Hassan HT, Zander A. Stem cell factor as a survival and growth factor in human normal and malignant hematopoiesis. Acta Haematol. 1996;95(3-4):257–262.

40. Nitsche A, Junghahn I, Thulke S, et al. Interleukin-3 promotes proliferation and differentiation of human hematopoietic stem cells but reduces their repopulation potential in NOD/SCID mice. Stem Cells. 2003;21(2):236–244.

41. Rasko JE, Metcalf D, Rossner MT, Begley CG, Nicola NA. The flt3/flk-2 ligand: receptor distribution and action on murine haemopoietic cell survival and proliferation. Leukemia. 1995;9(12):2058–2066.

42. Gilliland DG, Griffin JD. The roles of FLT3 in hematopoiesis and leukemia. Blood. 2002;100(5):1532–1542.

43. Rusten LS, Lyman SD, Veiby OP, Jacobsen SE. The FLT3 ligand is a direct and potent stimulator of the growth of primitive and committed human CD34+ bone marrow progenitor cells in vitro. Blood. 1996;87(4):1317–1325.

44. McKenna HJ, de Vries P, Brasel K, Lyman SD, Williams DE. Effect of flt3 ligand on the ex vivo expansion of human CD34+ hematopoietic progenitor cells. Blood. 1995;86(9):3413–3420.

45. Bernitz JM, Daniel MG, Fstkchyan YS, Moore K. Granulocyte colony-stimulating factor mobilizes dormant hematopoietic stem cells without proliferation in mice. Blood. 2017;129(14):1901–1912.

46. Tay J, Levesque JP, Winkler IG. Cellular players of hematopoietic stem cell mobilization in the bone marrow niche. Int J Hematol. 2017;105(2):129–140.

47. Jutila MA, Kurk S, Jackiw L, Knibbs RN, Stoolman LM. L-selectin serves as an E-selectin ligand on cultured human T lymphoblasts. J Immunol. 2002;169(4):1768–1773.

48. Shirure VS, Liu T, Delgadillo LF, et al. CD44 variant isoforms expressed by breast cancer cells are functional E-selectin ligands under flow conditions. Am J Physiol Cell Physiol. 2015;308(1):C68–78.

49. Hanley WD, Burdick MM, Konstantopoulos K, Sackstein R. CD44 on LS174T colon carcinoma cells possesses E-selectin ligand activity. Cancer Res. 2005;65(13):5812–5817.

