## Supplemental Material for "Knockout studies using CD34+ hematopoietic cells suggest that CD44 is a physiological human neutrophil E-selectin ligand"

**Contents:**

Supplemental Methods

Supplemental Movie A-C legends

Supplemental Tables S1-S3

Supplemental Figures S1-S5

**Supplemental Methods**

**Materials**: Directly conjugated mouse anti-human monoclonal antibodies (mAbs) were purchase from different suppliers. Anti-human Cutaneous Lymphocyte Antigen/CLA (HECA-452-eFluor® 660, CAT# 50-9857-82) and anti-human CD44 (Hermes, CAT# MA4400) were from ThermoFisher (Waltham, MA). MAbs against CD43 (MEM59- FITC, Cat# 315204), CD44 (BJ18-APC, Cat# 338805), CD33 (WM53-BV421, Cat# 303415) and CD16 (3G8- PerCP, Cat# 302029) were from BioLegend (San Diego, CA). Anti-CD15 (HI98- FITC, Cat# 555401) and CD11b (D12-PE, Cat# 347557) were from BD Biosciences (San Jose, CA). Polyclonal mouse anti-human IgG (H+L) (209-005-088) and peroxidase conjugated mouse anti-rat IgG (H+L) (212-035-082) were from Jackson ImmunoResearch (West Grove, PA). All recombinant selectin-human IgG1 fusion proteins were from R&D systems (Minneapolis, MN) and isotype control reagents were from BD Biosciences. All cytokines were purchased from Peprotech (Rocky Hill, NJ). All other reagents were from ThermoFisher or Sigma Chemicals (St. Louis, MO) unless otherwise mentioned.

**Cytospin:** 100µl cells at $10^6$/ml were added to CytoSep™ cytology funnels (Simport, Beloeil, Canada). Cells were centrifuged at 1000 rpm for 10 min in a Cyto-Tek Cyto centrifuge (Miles Scientific). The cells were then fixed using HARLECO® fixative solution I (Millipore-Sigma, St. Louis, MO), and stained using Hemacolor (Hemacolor®, EMD Millipore, St. Louis, MO). Images were acquired using Zeiss AxioObserver microscope using 40X objective/ 0.6 ph2 korr.

**RT-PCR:** Total RNA was isolated using the Trizol reagent. cDNA was generated by *in vitro* reverse transcription using primers designed against of the exon-exon junction of target genes (Primer-Blast tool; https://www.ncbi.nlm.nih.gov/tools/primer-blast/), and the cDNA synthesis kit (High-Capacity cDNA Reverse Transcription Kit). 30 cycles of PCR amplification were performed and the final product was resolved using 1% DNA agarose gel. Only one product with anticipated molecular mass was observed in each case. Primer pairs for each mRNA including the housekeeping gene (18sRNA) are listed in the **Supplemental Table S1**.

**Flow cytometry:** 1-10µg/mL fluorescently labeled mAbs were incubated with $2\times10^6$cells/mL in HEPES (30mM 4-(2-hydroxyethyl)-1-piperazineethanesulfonic acid, 110mM NaCl, 10mM KCl, 1mM MgCl2, 10mM Glucose, pH 7.4) containing 0.1% human serum albumin (HSA) and 1.5mM $CaCl_2$ for 20min at 4ºC. The cells were then washed, resuspended and analyzed using a Fortessa X-20 flow cytometer (BD Biosciences). For the selectin-IgG binding assay under static condition, 3-10 µg/mL recombinant human P-/E-/L-selectin-IgG fusion proteins were complexed with 10µg/mL PerCP conjugated goat anti-human IgG Ab for 10 min at RT as described previously [1]. This mixture was then added to a 20µl leukocyte suspension containing $2\times10^6$ cells/mL. Following a 10 min incubation at RT, the samples were analyzed using flow cytometry. For cell viability analysis, 1µL of Annexin-V FITC was mixed with 10µg/mL LDS-751 in HEPES buffer containing 10mM calcium, and added to 5µL cells at $10^6$/mL. Following 10 min incubation at 37°C, the sample was diluted 50-fold and read using a flow cytometer.

***In vitro* transcription to produce sgRNA**: To make native sgRNA**,** a 54-56bp custom DNA oligo was purchased that contained the 18bp T7 promoter followed by a 19-21bp guide sequence against the target gene based on published literature [2,3], and the first 17 bases of the sgRNA scaffold (Supplemental Table S2). To produce the corresponding synthetic sgRNA, we used the EnGen sgRNA Synthesis Kit (New England Biolabs, Ipswich, MA). To this end, the above oligonucleotide was mixed with T7 enzyme and reaction mix provided with the kit for 2 h at 37ºC. The product formed was DNAase treated, and the RNA was purified using the RNA Clean & Concentrator kit (Zymo Research, Irvine, CA). The final product was stored at -80ºC until use. All

**Western blot analysis of CD44-Fc and CD44-FUT7-Fc**: In the western blot studies, 30µl of cell culture supernatant was loaded into Novex$^{TM}$ WedgeWell 4-20% Tris-glycine gels (Thermofisher) along with mock control in a mini gel tank (Thermofisher). The condition for gel running was 90V 30min, followed by 120V 60min. Transfer was done by using Trans-Blot Turbo Transfer System (Biorad, Mississauga, ON). Proteins on the blots were then detected using 0.5µg/mL anti-CD44 Hermes-1 or anti CLA (HECA-452), along with HRP conjugated secondary antibodies. All data were acquired using ChemiDoc MP Imaging system (Biorad).

**Flow cytometry quantitation of CD44-Fc and CD44-FUT7-Fc**: In the flow cytometry studies, mouse anti-human polyclonal antibodies (Jackson ImmunoResearch) were coupled to Polybead® Carboxylate Microspheres (Polysciences Inc, Warrington, PA) using carbodiimide chemistry. To this end, ~$80x10^6$ beads were washed and then activated by addition of 0.05M sulfo-NHS (N-hydroxysulfosuccinimide, Thermofisher) along with 0.05M EDC (1-ethyl-3-(3-dimethylamino) propyl carbodiimide, hydrochloride; G-Bioscience, St. Louis, MO), overnight at RT on a vortex. The next day, the activated beads were washed several times with PBS (phosphate-buffered saline) and resuspended in 100µl volume at $10^8$ beads/mL. ~20-40 µg polyclonal antibody was then added to these beads for coupling for 2-3h at RT with mixing. Next, the beads were washed with PBS and unreacted sites quenched with 40mM Ethanolamine (Alfa Aesar, Ward Hill, MA) in ~100µL volume for 30 min at RT. The bead surface was then additionally blocked for non-specific binding by resuspending beads in PBS containing 1% BSA. To detect the CD44-Fc fusion protein, 1mL of culture supernatant was incubated with $1x10^6$ goat anti-human polystyrene beads prepared as described on a shaker for 30min at RT. After two PBS washes, bead-bound CD44-Fc was detected using 2µg/mL fluorescent tagged anti-human CD44 mAb using standard flow cytometry.

**References (also cited in main manuscript)**:


1. Stolfa G, Mondal N, Zhu Y, Yu X, Buffone A, Jr., Neelamegham S. Using CRISPR-Cas9 to quantify the contributions of O-glycans, N-glycans and Glycosphingolipids to human leukocyte-endothelium adhesion. *Sci Rep*. 2016;6:30392.
2. Tzelepis K, Koike-Yusa H, De Braekeleer E, et al. A CRISPR Dropout Screen Identifies Genetic Vulnerabilities and Therapeutic Targets in Acute Myeloid Leukemia. *Cell Rep*. 2016;17(4):1193-1205.
3. Wang T, Birsoy K, Hughes NW, et al. Identification and characterization of essential genes in the human genome. *Science*. 2015;350(6264):1096-1101.

**Supplemental Movie legends:**

***Movie A: Preventing secondary adhesion***. Neutrophils derived from hHSCs were rolled on E-selectin substrates at 1 dyn/cm$^2$. Clusters of cells were observed in the absence of blocking mAb (Top panel). These clusters were abolished upon addition of either anti-L-selectin mAb DREG56 (Middle panel) or anti-PSGL-1 mAb KPL-1 (Bottom panel). Thus, these clusters are due to secondary L-selectin PSGL-1 dependent neutrophil-neutrophil interactions.

***Movie B: Confocal microscopy rolling assays with neutrophils derived from CD34+ hHSCs.*** Microfluidics based cell adhesion studies were performed with a mixed neutrophil population including single and dual CD43/CD44 KOs at 1dyn/cm$^2$. All cells were labeled with CD43-FITC and CD44-APC. Top panel: FITC channel data showing cells expressing CD43. Middle panel: APC channel showing cells expressing CD44. Bottom panel: Bright-field plus fluorescence merged channel showing CD43/CD44 DKOs (white cells), CD44-SKOs (green), CD43-SKOs (red), and unmodified/wild-type cells (yellow). Data from all channels were individually analyzed to characterize the phenotype of the individual rolling cells.

***Movie C: Rolling phenotype of CD44-Fc bearing microspheres on E-selectin***. Top panel: Robust rolling of beads coupled with CD44-FUT7-Fc at a wall shear stress of 1dyn/cm$^2$. Bottom panel: Beads bearing CD44-Fc do not roll on E-selectin substrates.

**Table S1: RT-PCR primer list**

| Gene name | Forward (5' to 3') | Reverse (5' to 3') |
|---|---|---|
| 18sRNA | CAGCCACCCGAGATTGAGCA | TAGTAGCGACGGGCGGTGTG |
| MMP9 | CGCTGGGCTTAGATCATTCCT | TTCAGGGCGAGGACCATAGA |
| MPO | TGCTGGAAGGTGGCATTGAC | CATGTTCAGAGCAGGCAGGT |
| ELANE | CACTGCGTGGCGAATGTAAAC | GACCCGTTGAGCTGGAGAATC |

**Table S2: sgRNA sequence for gene editing***

| Gene name | sgRNA sequence (5' to 3') |
|---|---|
| CD43 | CACCAATGGAAGTCCAAAG |
| CD44 | GTGCCACCAAAACTTGTCCA |

* For *in vitro* transcription, the template DNA included T7promoter-$(N)_{20}$-17 bases of the sgRNA scaffold, i.e. TTCTAATACGACTCACTATA-$(N)_{20}$-GTTTTAGAGCTAGA. $(N)_{20}$ is listed in the table above.
* For synthetic RNA, the first three bases were protected using 2'-O-methyl 3' phosphorothioate inter-nucleotide linkages. This is indicated by * for the case of CD44: G*U*G*CCACCAAAACUUGUCCA+synthego modified EZ scaffold

**Table S3: PCR primers for cloning CD44***

| | |
|---|---|
| Forward primer | agatccGCTAGCATGGACAAGTTCTGGTGGCAC |
| Reverse primer | aagctaTCTAGATTTGTCATCGTCATCCTTGTAGTCGCCAGGGATCTGGGGGGTTC |

* Lowercase indicates handles for restriction enzyme digestion; green: Restriction enzyme site; Blue: Enterokinase cleavage site added after CD44.

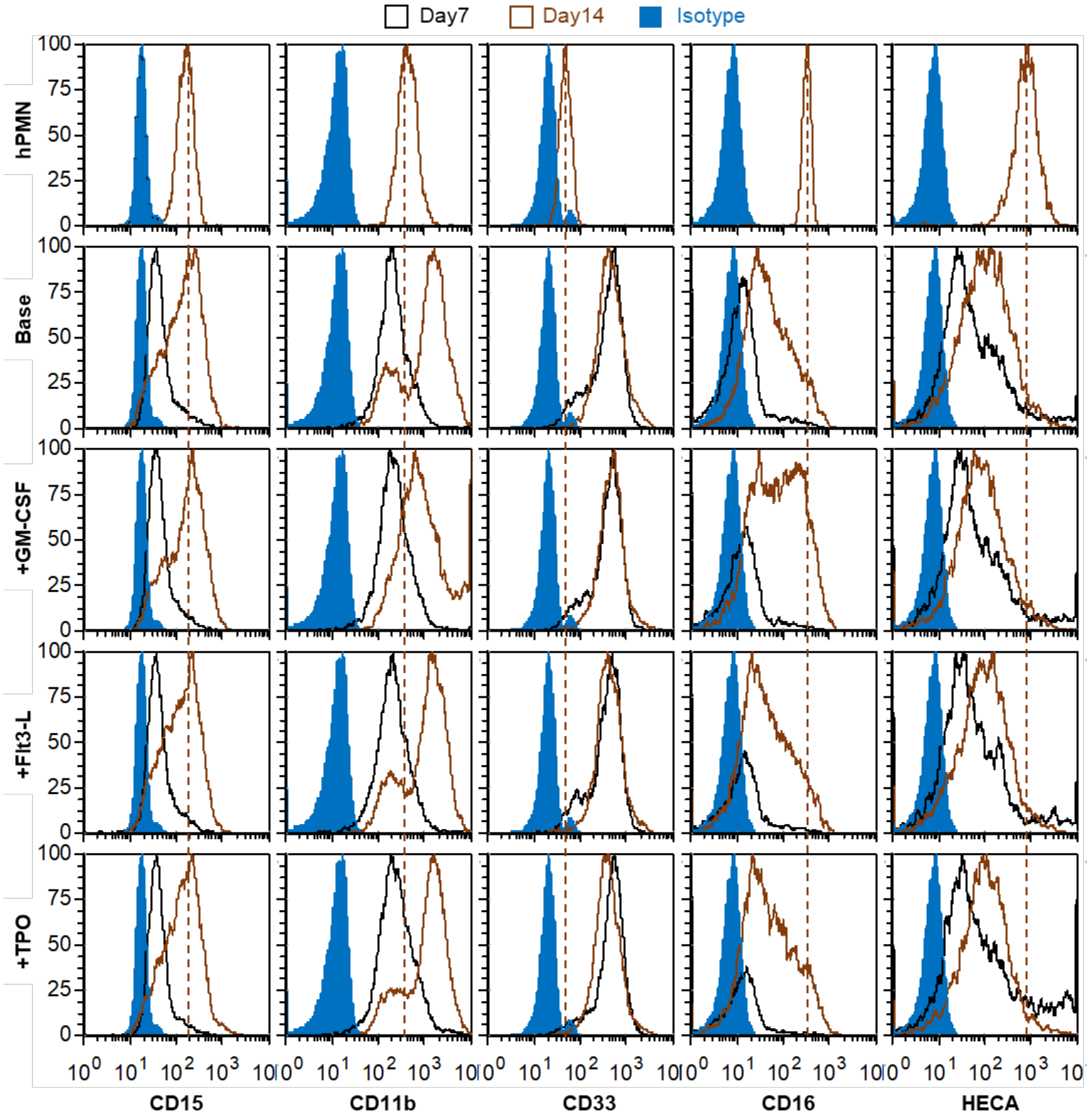


**Supplemental Figure S1. Epitope expression on neutrophils differentiated from hHSCs**. Flow cytometry was used to measure myeloid markers. Compared to peripheral blood neutrophils (dashed vertical line), all cells differentiated using different chemokine cocktails displayed increased CD11b and CD33 expression, but reduced CD16 levels. CD15 levels were comparable in all cases to primary neutrophils. The sLe$^{X}$ binding HECA-452 epitope was lower at day 7, but its levels markedly increased during the late stages of the differentiation process between days 7-14. Data are representative of >4 repeats for each condition.

**Zhu et al.**
**Figure S2**

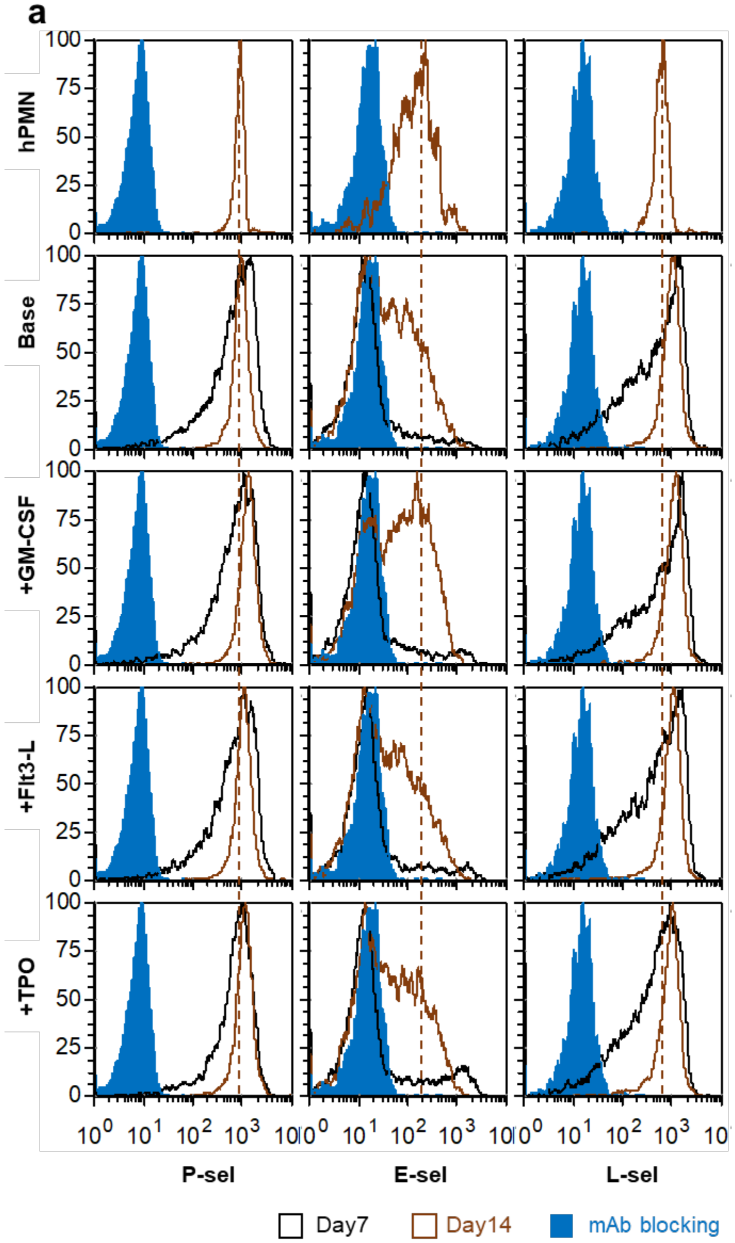


**Supplemental Figure S2. Binding of recombinant selectin IgG to human blood neutrophils and differentiated cells**. The extent of L- and P-selectin IgG binding to differentiated cells was comparable to human peripheral blood neutrophils both at day 7 and day 14. E-selectin Ig binding was however low at day 7 and different cell sub-populations were observed. At least a portion of these cells exhibited robust E-selectin binding comparable to peripheral blood neutrophils at day 14. Blocking antibodies (against L-/P-/E-selectin) confirm the specificity of the measured binding interactions. Data are representative of >4 repeats for each condition.

Zhu et al.
Figure S3

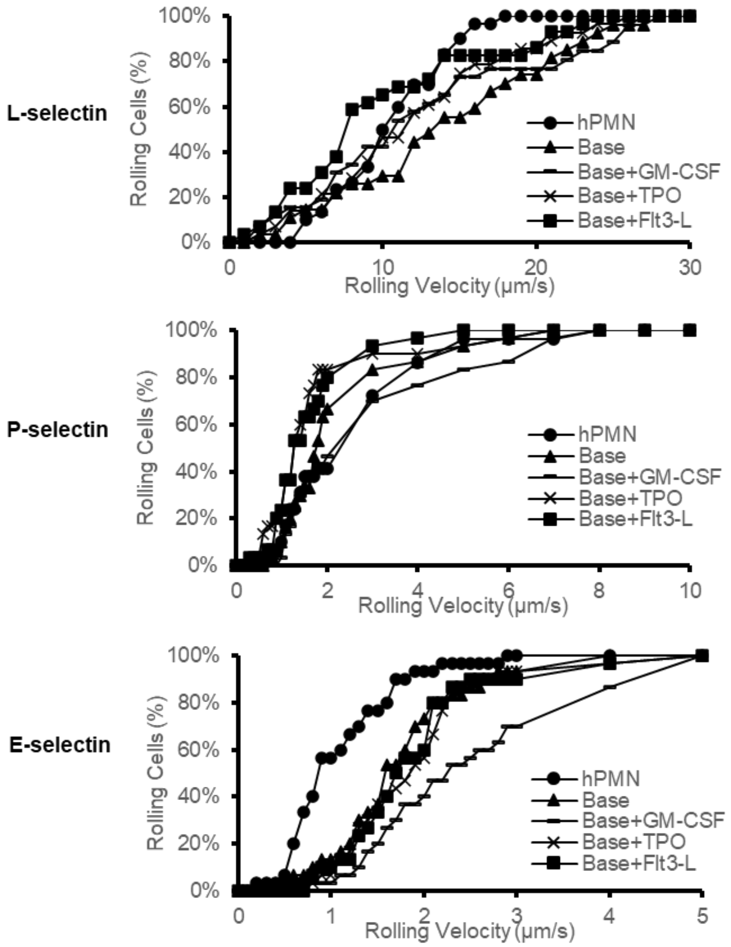


**Supplemental Figure S3**. **Rolling velocity of differentiated neutrophils vs peripheral blood neutrophils**. The rolling phenotypes of all cell types was similar on substrates bearing L-selectin (top panel) and P-selectin (middle panel). The rolling velocity of peripheral blood neutrophils was lower on E-selectin substrates, compared to neutrophil-like cells differentiated from CD34+ hHSCs (bottom panel). Wall shear stress = 1 dyn/cm$^2$.

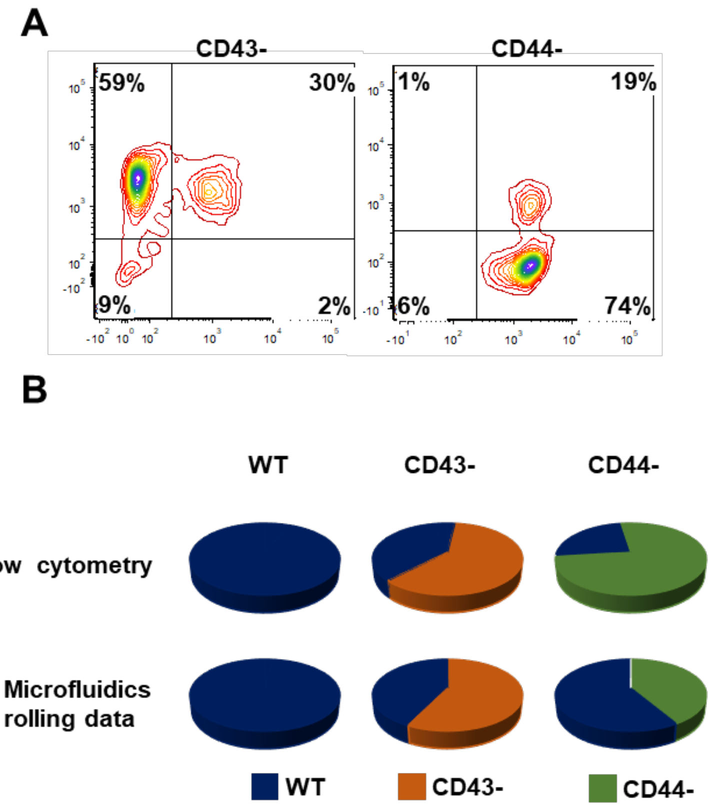


**Supplemental Figure S4**. **Knock efficiency for CD43- and CD44- single KOs. A**. RNPs targeting CD43 and CD44 were produced by mixing 1µg Cas9, 1µg CD43 modified sgRNA or 1µg CD44 modified sgRNA in 2µL volume. These were electroporated into hHSCs at day 2 ($0.1\text{-}0.25\times10^6$ cells) followed by differentiation to neutrophils. Flow cytometry measured receptor expression at day 14. Knockout efficiency was 68% and 80% for the single CD43- and CD44- knockouts. **B.** Cells labelled with anti CD43-FITC and anti CD44-APC were perfused over E-selectin substrates at 1 dyn/$cm^2$. Cell rolling density was measured using confocal microscopy with antibody labels being used to distinguish wild-type cells from individual knockouts. Data are presented in the form of pie chart, in order to compare the editing efficiency of the different cell populations using flow cytometry (top charts) vs. the fraction of rolling cells observed in the confocal flow-shear assay (bottom charts). Results show that CD44 is an important neutrophil E-selectin ligand.

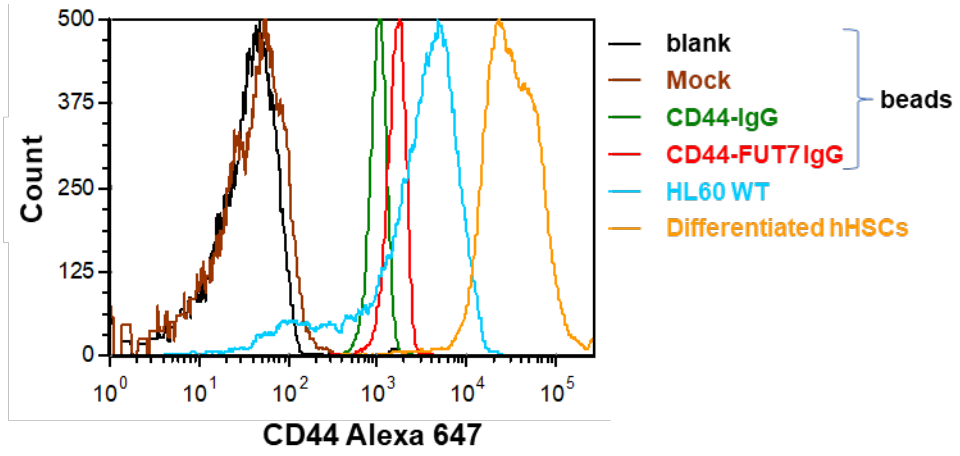


**Supplemental Figure S5**. **CD44 expression on different beads and cell types.** Fluorescent anti-CD44 antibody was used to detect CD44 expression using flow cytometry on i) beads coupled with mock transfection medium, CD44-FUT7-Fc and CD44-Fc, ii) wild-type HL60, and iii) hHSCs differentiated to neutrophils. Data are representative of 4-6 independent repeats.
